# Synthesis of Metal-Organic Framework (MOF) Based Composites for pH-Responsive Drug Delivery

**DOI:** 10.1101/2025.06.18.660385

**Authors:** Aayushi Kunde, Dhiraj Bhatia

## Abstract

Metal-Organic Frameworks (MOFs) are nanoporous, and nanostructured resources crafted from hydrocarbon-based linkers connecting inorganic metal nodes. A considerable amount of curcumin can be encapsulated within MOF’s porous structure, shielding it from deterioration. The curcumin’s solubility was significantly improved by including carbon quantum dots (QDs) into the nanocomposite. In this study, curcumin was encapsulated using the double emulsion approach using MOF-based nanoparticles and pH-sensitive chitosan/PEG hydrogel. The nanocarrier was characterised by X-Ray diffraction (XRD), Fourier transform infrared spectroscopy (FTIR), field emission scanning electron microscopy (FE-SEM), Dynamic light scattering (DLS), and zeta potential. UV spectroscopy and spectrofluorimetric provided insights into the binding of curcumin to the added QDs. Curcumin loading and encapsulation efficiency were found to be about 31% and 85%, respectively. The release profile of curcumin showed pH-dependence and controlled release by the nanocomposite, following the K-P diffusion model. Biocompatibility against the MCF-7 breast cancer cells validated the anti-cancer potential of the nanocarrier.

## 1. Introduction

An enormous number of people suffer from cancer each year, and it is considered among the deadliest. Numerous approaches to combat cancer have been investigated. One way to treat cancer is with chemotherapy; however, it comes with many other limitations. The increasing popularity of conventional medicine, especially plant-based, has sparked a lot of studies on the therapeutic value of compounds with natural origins. It has been established that curcumin affects cancer cells by disrupting the cycle of cell division. However, the hydrophobic characteristic of curcumin hinders its ability to pass through cellular membranes, as it forms H-bonding and shows hydrophobic relationships with the fatty acyl chains that constitute the membranes. This has caused the curcumin concentrations to stay minimum in the cytoplasm. Curcumin was found to have less potential due to its pharmacological disadvantages, which include reduced bioavailability and water insolubility. Consequently, research has proposed several strategies to address these issues, such as the application of derivatives of curcumin or nano-based systems for delivering drugs^1–13^. The nanodrug delivery system for delivering curcumin can bypass these obstacles by increasing its solubility, bioavailability, and targeted delivery.^14–20^ This study aims to investigate a workable plan to synthesise a pH-sensitive nano-composite as a nanocarrier for enhancing curcumin’s transport and efficacy in cancer treatment.

Approaches are currently being explored to minimise immunogenicity by coating or chemically functionalising the nanostructures with various substances, including polymers, natural biomolecules, cell membranes, configurable surfactants, peptides, etc., while also improving the specificity for targeted delivery. Effective drug-loaded nanocarriers, known as nano-drug delivery systems (DDS), consist of biologically compatible and biodegradable materials, including metals, lipids, and synthetic and natural polymers. These materials typically range in less than one thousand nm in diameter^10,14–19,21–44^.

Numerous categories of coordination polymers have grown exponentially in the last few decades, and MOFs have become more well-liked than other systems because of their improved biocompatibility and adaptable loading capacities^45–49^. When it comes to drug loading into MOFs, there are two main approaches. The first is the impregnation approach, in which medications are inserted into MOF pores via capillary forces, coordination processes, or electrostatic interactions. An in-situ encapsulation during the MOF development process, using a one-pot technique, can integrate functional molecules and build drug/MOF composites. Another approach is to wrap the outermost layer of MOFs with different polymers. Chitosan addition to MOF may improve MOFs’ dispersion, provide chemical stability and bio-adhesive qualities, besides decreasing undesirable immunological reactions^23,45–50^.

With a particle size of less than 10 nm, carbon quantum dots (QDs) represent a novel family of carbon nanostructures. Small carbon clusters enhanced by nitrogen, oxygen, and hydrogen, known as carbon quantum dots, are useful for bioimaging and drug administration because of their water solubility and potent luminescence phenomena^38,51–54^. The ability of their functional groups, such as –NH_2_, -CH_2_-OH, and -C-O-C-, to interact with those of other biomolecules is intriguing for the construction of new nanocomposites with enhanced characteristics ^7,8,52,55^. Curcuminoids and carbon dots are used as essential components of the drug cargo rather than just as carriers or therapeutics ^5,6,56^. The new approach exploits the acidic tumour microenvironment to attain selective drug release by incorporating pH-sensitive ester connections. In this study, a double emulsion method was used to generate a membrane around the nanodrug that led to better drug retention and the formation of spherical nanocarriers with a smaller particle size. The oil layer that separates the two water layers controls the drug release rate. The presence of multiple layers in the double emulsion system can increase the release time and develop a slow and sustained release profile for curcumin, which is of paramount importance for drugs that have a low bioavailability.

## 2. Experimental

### 2.1 Materials and Methods

PEG 400 was provided by SRL, and acetic acid (glacial) was procured from Merck. SPAN 80 (≥60 %), curcumin (CU) (C_21_H_20_O_6_), and imidazolate-2-carboxyaldehyde (ICA), were provided by Sigma Aldrich. Zn(NO_3_)_2_•6H_2_O, Trioctylamine (TOA), and Chitosan (HMW, DA > 90 %) were provided by TCI chemicals. MTT was procured from Sigma Aldrich. Olive oil was purchased from a local supplier. MCF-7 cell line, 0.25% Trypsin-EDTA, FBS, was acquired from Gibco.

### 2.2 Preparation of precursors

#### 2.2.1 ZIF-90, a metal-organic framework (ZIF)

The reported procedure was modified for the synthesis of ZIF-90^57^. The procedure involves the dissolution of ICA and Zn salts separately in a 1:1 ratio. Solution A was prepared by mixing 20 millilitres of DMF with zinc acetate (Zn(CH_3_COO)_2_·2 H_2_O). For solution B, 5 millilitres of trioctylamine were mixed with the required amount of imidazole-2-carboxaldehyde (ICA). Add solution B to solution A gradually while stirring constantly. Stir the mixture while heating it to a predetermined temperature (60°C) for two hours. The precipitate is collected by centrifuging the mixture, and ethanol is then used to wash it. Dry the obtained solid material in an oven at 100°C for 24 hrs.

**Scheme 1.**
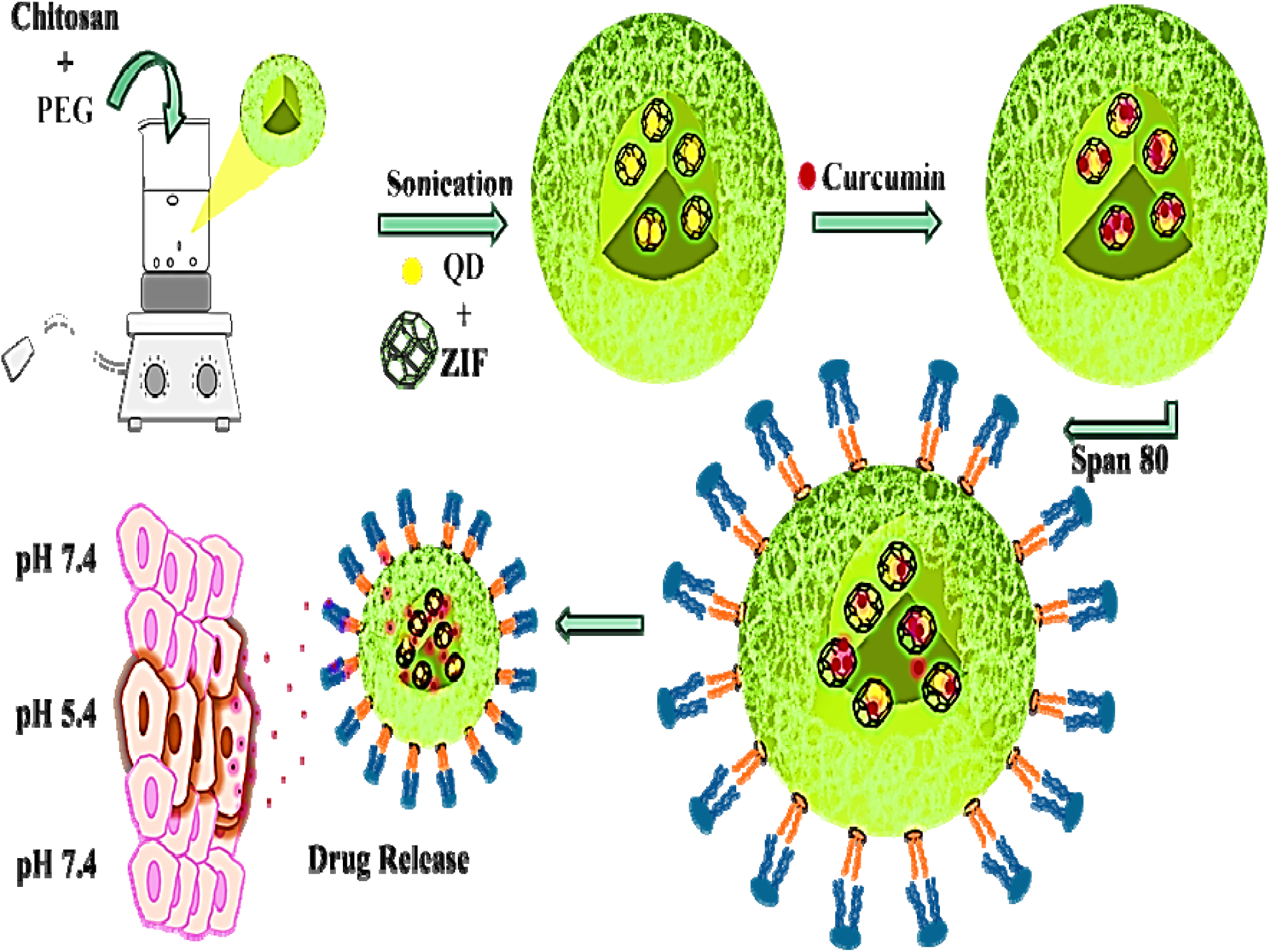
Graphical representation depicting the synthesis of curcumin-loaded DDS.

#### 2.2.2 Preparation of carbon quantum dots (NCQD)

The hydrothermal procedure for the preparation of quantum dots (NCQD) was similar to the reported procedure with little modification. The mixture of citric acid (0.6g) and urea (1.0 g) was dissolved in 20 ml of distilled water and then heated in a stainless-steel autoclave at 200 °C for 12 h^58^. The brown solution obtained was purified and freeze-dried. The double emulsion approach was followed to generate drug delivery systems for encapsulation of both water-attracting and water-repelling drug molecules^59^.

#### 2.2.3 Preparation of curcumin-loaded drug delivery system (DDS)

For encapsulating curcumin in the nanocarrier, a double emulsion method was utilised. 2% Chitosan-PEG mixture (70:30 w/w) was made with a 5% acetic acid solution under stirring at room temperature. 5 mg/ml of NCQD and 10 mg/ml of ZIF-90 were mixed in the above polymer hydrogel to get a homogeneous mass. After adding the required quantity of curcumin, 0.2% (v/v) Span 80, and 15 ml of olive oil were added to the hydrogel. Finally, 15 mL of distilled water was added to the above-prepared hydrogel and left to attain a steady condition. After separating the oil phase by a sampler, the curcumin-loaded nanocarrier was separated from the remaining solution by centrifugation for 5 min at 6000 RPM. Different composites, namely Chitosan/PEG, Chitosan/PEG/NCQD/ZIF, and Chitosan/PEG/NCQD/ZIF/Curcumin at every stage, were separated for freeze drying to study the effect of precursors.

### 2.3 Loading and encapsulation efficiency

A pre-weighted amount of freeze-dried composite was dispersed using one ml of PBS to find the efficiency of the prepared system. To phase out the ethyl acetate layer, a known amount of ethyl acetate was introduced to the solution containing the dispersed sample. Once the two layers get separated, the amount of curcumin present in the ethyl acetate phase was calculated using equations (2.1) and (2.2) and expressed as loading efficiency and curcumin encapsulation percentage^60^. Experiments were repeated twice for accuracy.

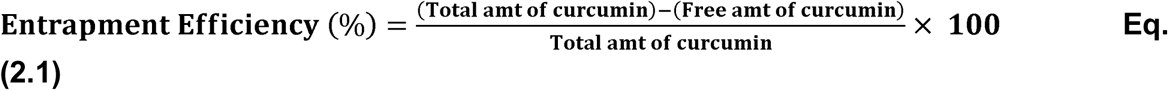

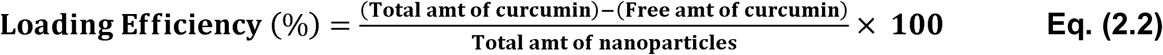

### 2.4 Optical Properties

Encasing QDs in a nanocomposite provided a novel luminous nanoplatform. At 365 nm, the aqueous solution containing NCQD exhibits brilliant blue fluorescence that is easily visible to the unaided eye. A decrease in fluorescence intensity brought on by one of a number of processes, including collisional quenching, reactions during excitation, energy transfer, and complex-forming ability, is referred to as fluorescence quenching. The interaction between a quencher molecule and a fluorophore is the primary form of quenching in sensing. The fluorophore and quencher collide either dynamically or statically to cause this occurrence. A static-based quenching process frequently takes place between the two species, and the majority of bioassays use a probe that consists of a fluorophore and quencher. When the quencher and fluorophore combine to form a complex in the ground state, the fluorophore is unable to absorb light and move an electron into the excited state, a phenomenon known as static-based quenching. The fluorescence returns to the lowest ground level as a result of collisions between the quenching molecule and the excited fluorophore in dynamic-based quenching.

### 2.5 Evaluation of curcumin release

Using the dialysis approach, the discharge of curcumin from the composite was examined in two distinct PBS solutions. First, a dialysis tubing (MWCO = 1.2 kDa) was filled with 1 ml of composite emulsion. The tubing was then submerged in 100 millilitres of PBS and kept in a water bath under continuous stirring at 37 °C. One millilitre of the sample was separated after the required time and replaced with a similar volume of PBS buffer at predesigned intervals from 0 to 96 hours. Curcumin’s calibration curve provides the amount of drug released in PBS at 425 nm directly. The cumulative medication release % was then calculated using Eq. (2.3)^61^. The experiment was repeated while maintaining pH values of 7.4 and 5.4.

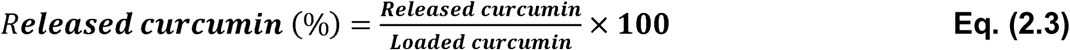

### 2.6 Kinetics of curcumin release

For studying the process of curcumin released from the nanocomposite at 7.4 and 5.4 pH, the concentration values of curcumin released were applied to a number of models and the best-fitted model was identified using R^2^, which shows the goodness of fit. Different models that were used for fitting data are listed below:

## Zero order

The zero-order model would be used to fit the drug discharge kinetics of optimal DDS with constant drug levels during the drug release. One way to depict the **zero-order** release is given as^62^:

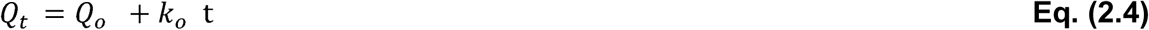

## First order

The first-order model is a semi-empirical kinetic model^63^ that is frequently used to forecast drug release from porous materials, mostly for H_2_O-soluble medications. It can be expressed mathematically as:

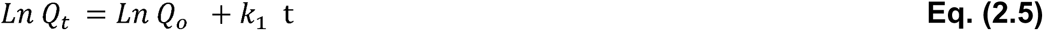

In both these models, Q0 and Qt are the amount of the drug released as the time varies from zero to time t, whereas k_1_ is the discharge constant with respect to order.

## Korsmeyer–Peppas model

A comprehensive yet straightforward semi-empirical equation of the Korsmeyer–Peppas model aids in simulating many processes that contribute to the release rate or unknown release mechanisms. One way to depict Korsmeyer-Peppas release is as follows:

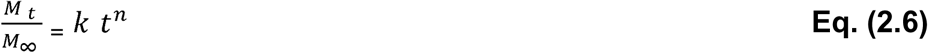

where n and k are the release exponent and constant, respectively. Mt/M∞ signifies the medication released fraction at time ‘t’. The most frequent method of drug release when n is less than 1/2 is Fickian diffusion, in which the rate of release is determined by solvent penetration. Non-Fickian processes control the rate of drug release, and diffusion alone does not control the release; swelling may also be involved, with n greater than half but less than one^64^.

## Higuchi model

Both water-attracting and water-repelling substances that are released from drug delivery systems using this model, as per the equation shown below:

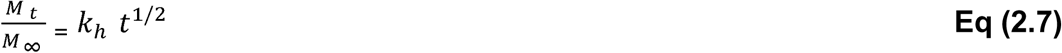

where k_h_ is the Higuchi release constant^62^.

### 2.7 MTT Assay

To assess the efficacy of the curcumin-loaded Chitosan/PEG/NCQD/ZIF in gel form, 5 × 10^3^ MCF-7 cells were seeded into a 96-well plate, and then the cells were treated for 72 hours with (1-200 µg/mL) working doses made in complete media. 0.5 mg/ml of working MTT solution was added after treatment, and the mixture was then incubated for three hours. After dissolving the resultant formazan crystals in DMSO, they were left to stand for fifteen minutes. Absorbance at 570 nm was measured using the Byonoy® 96 absorbance plate reader. The percentage of cells that survived to control (without the medicine, only complete media) was calculated and plotted against different drug concentrations when n = 3 using GraphPad Prism 8 software. Following ten minutes of plate shaking, the optical absorbance of the solution at 570 nm was measured, and the cell viability was calculated using Eq. (2.8)^65^.

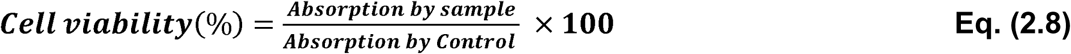

### 2.8 Characterization techniques

#### 2.8.1 PXRD

This method identifies different phases of the compound in powder form after synthesis. The phase of the material can be compared to any previously published crystal structure from a database by using the 2θ angles or the d-spacing with the relative intensities. The crystalline quality of the nanocomposite was inspected by XRD between 10° and 80°.

#### 2.8.2 FESEM

This method depicts the morphologies of the synthesised composite at different magnifications and an acceleration voltage of 20 kV.

#### 2.8.3 FTIR

This is a characterization tool that assesses a sample’s absorption of IR light to identify chemical compounds and analyse molecular structures. The purpose of this analysis was to examine the functional groups and chemical makeup of room-temperature nanocomposites. Samples ranging from 400 to 4000 cm^−1^ were subjected to FTIR analysis for the intermediates and final composite.

#### 2.8.4 DLS and Zeta analyser

The particle size and its distribution in a suspension can be examined using this method. It gauges variations in the strength of dispersed light, which are connected to the particles’ Brownian movement. The nanocomposite’s particle size and surface charge at 25 °C are then ascertained using this data. The samples were sonicated for fifteen minutes to quantify the zeta potential, and the analyses were carried out at a 90° scattering angle.

## 3. Results and Discussion

### 3.1 FT-IR Spectroscopy

The spectra of the as-synthesised composites and the precursors used in their synthesis are shown in Fig.1 (A, B, C). In Fig. 1A, the spectra of PEG, Chitosan, and the Chitosan-PEG composite are shown. O-H and N-H stretching show a large peak at 3347 *cm*^−l^ , which suggests the existence of intermolecular hydrogen bonds^66^. Stretching vibrations between carbon and hydrogen are accountable for the smaller bands seen at 2862 cm^−1^. Furthermore, the *> c = o* and *c-N* ν(s) of the *-coNH*_2_ I & III bands are linked to the maxima at about 1600 and 1369 *cm*^−l^ , respectively [19]. In the PEG spectra, the bands seen at 2884 & 3464 *cm*^−l^ represent γ(s) and 1473 and 1345 *cm*^−l^represent the δ(b) of alkyl and hydroxyl groups, respectively. The C-O-H ν(s) are associated to the 1080–1160 *cm*^−l^range^67^.The synthesized composite of chitosan and PEG shows similar spectra to those observed in the individual band structures, but the broadening illustrates hydrogen bond formation amongst the *-oH* grp of PEG and the *-oH & NH*_2_ group of chitosan^68^. Functional groups in CNQD and MOF were also characterized (Fig. 1B). In the ZIF-90 Spectra the peaks between 1200-1400 *cm*^−l^and the peak around 580 *cm*^−l^indicate the presence of imidazole moieties of the MOF used, the band at around 1685 *cm*^−l^ depicted >C*=o* ν(s) of RCHO groups^69,70^. The synthetic material did not take part in the reaction, as evidenced by the fact that it’s C=O bond’s peak position remained unchanged. Imidazole-2-carboxaldehyde’s N underwent a coordination process with Zn^2+^, as evidenced by the strong peak at 1200–1400 *cm*^−l^and the preservation of the aldehyde group’s distinctive peak at 2700–2900 cm^−1^. The ν(as) & ν(s) of C–O–C functional groups, which are classified as ether or epoxy, are represented by the peaks seen at 1321 and 1194 respectively^71^. Broadband around the region 3204 cm-1 -3405 corresponds to hydroxyl and amine ν(s)^72^. These results indicate that the synthesized NCQDs had sufficient hydrophilic groups for attaining stability in water^73^.

**Fig. 1.**
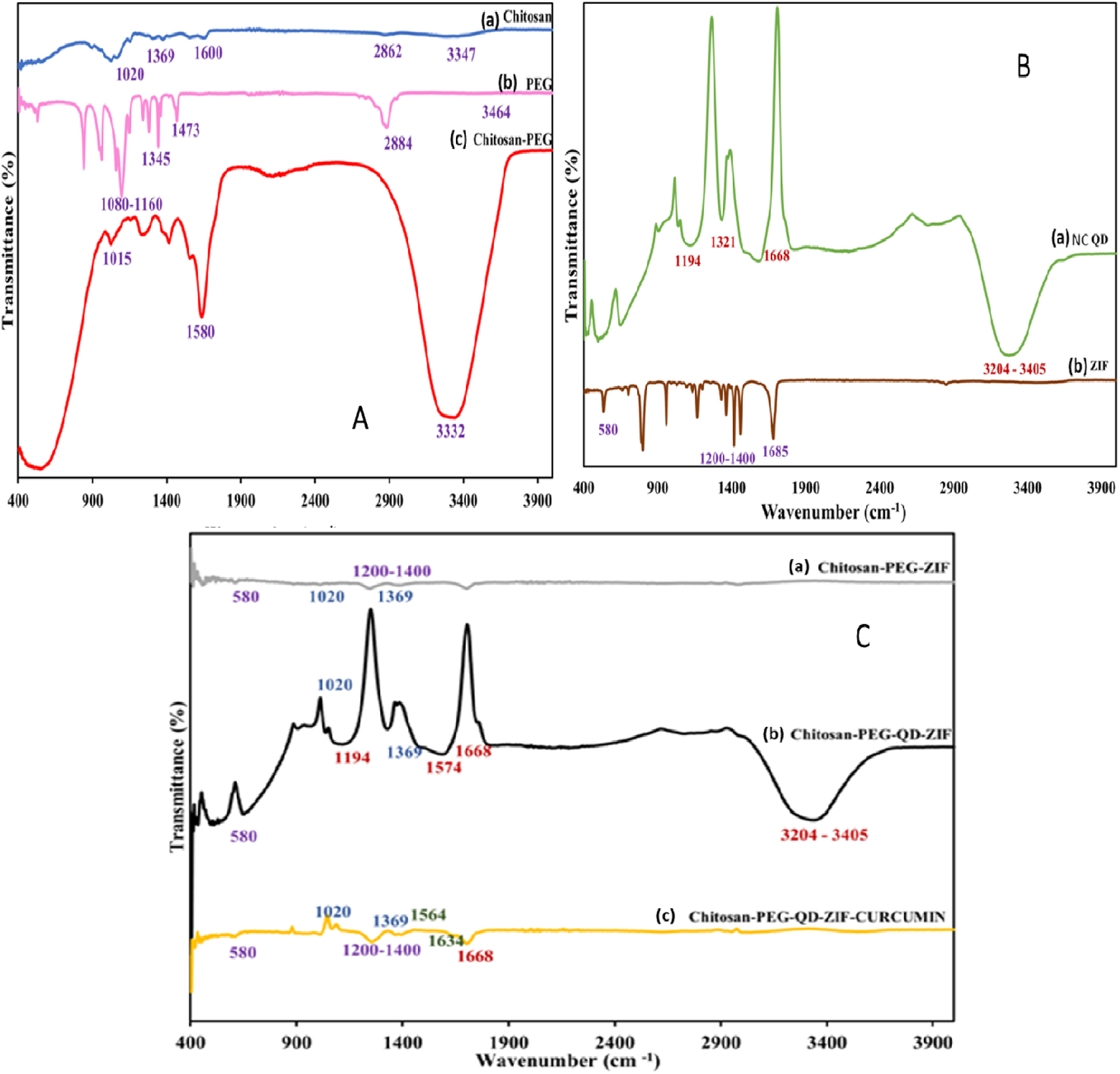
FTIR of A: Chitosan (a). PEG (b), Chitosan/PEG (c); B: QD (a), ZIF (b); and C: Chitosan/PEG/ZIF (a), Chitosan/PEG/QD/ZIF (b) Chitosan/PEG/QD/ZIF/CU (c)

A drift in the peak intensity ∼1668 *cm*^−l^ as can be seen in Fig. 1B due to overlapping of *>c = o*in NCQD and *N-H (amide II)* bands in chitosan. All the results are well aligned in the final composites’ resulting spectrum as previously established in individual spectra. In Fig. 1C the spectra of curcumin-loaded composite show smaller bands at 1564 and 1634 *cm*^−l^ that signifies *c= c* and *c= o* ν(s) bonds of curcumin^66^. When curcumin was added to the composite, all of the distinctive peaks of Ch/PEG/NCQDs were still present. Curcumin and Ch/PEG interaction is indicated by a noticeable fall in intensity for peaks at 1580 and 1015 *cm*^−l^. The emergence of the same distinctive peak at 1668 *cm*^−l^ in the composite that appeared in NCQDs spectrum, indicates that the – *cooH* is present on the surface of NCQDs coordinates with Zn^2+^ and functions as an “antenna” for sensitising the luminescence. However, it is seen that the distinctive peaks’ strength diminished compared to that of pure NCQD, which could be because NCQDs have been successfully incorporated into the composite. Through electrostatic interactions amongst the carboxyl and hydroxyl groups of NCQDs and the *-oH* and *R-o-R* groups of the hydrogel, curcumin forms a bond with the surface of CQDs. The *-cooH* and *-oH* groups on the surface of CQDs have the ability to bind via H bonding. The amine group gets protonated at acidic pH. The carboxyl groups are also impacted by this protonation, which alters the electrostatic connection and has a big impact on drug release.

### 3.2 XRD

Fig. 2 investigates the crystallographic structures of the synthesized composites, including intermediates. The sharp and intense peaks of ZIF-90 in Fig.1 (a) show a highly crystalline structure. A broad peak at 2θ = 19° in Fig.1(b) confirms the amorphous character of chitosan, also depicting characteristic peaks of PEG at around 2θ = 19.2° and 23.3°, consistent with the literature^74^. No shift in peak position was observed in Fig.1 (b, c), indicating a stable crystal structure formation leading to successful incorporation of every component in the matrix of Chitosan/PEG and Chitosan/PEG/ZIF. The XRD pattern for curcumin-loaded composite in Fig. 1(d) reveals diffraction peaks at 19.2° and 23.3°. The peaks show a slight increase in intensity and a bit of broadening, suggesting that the drug has been effectively incorporated into the nanocarrier. These findings are consistent with earlier research on polysaccharide-based nanocarriers intended for the delivery of curcumin, which found that drug loading somewhat decreased the degree of crystallinity^75^.

**Fig. 2.**
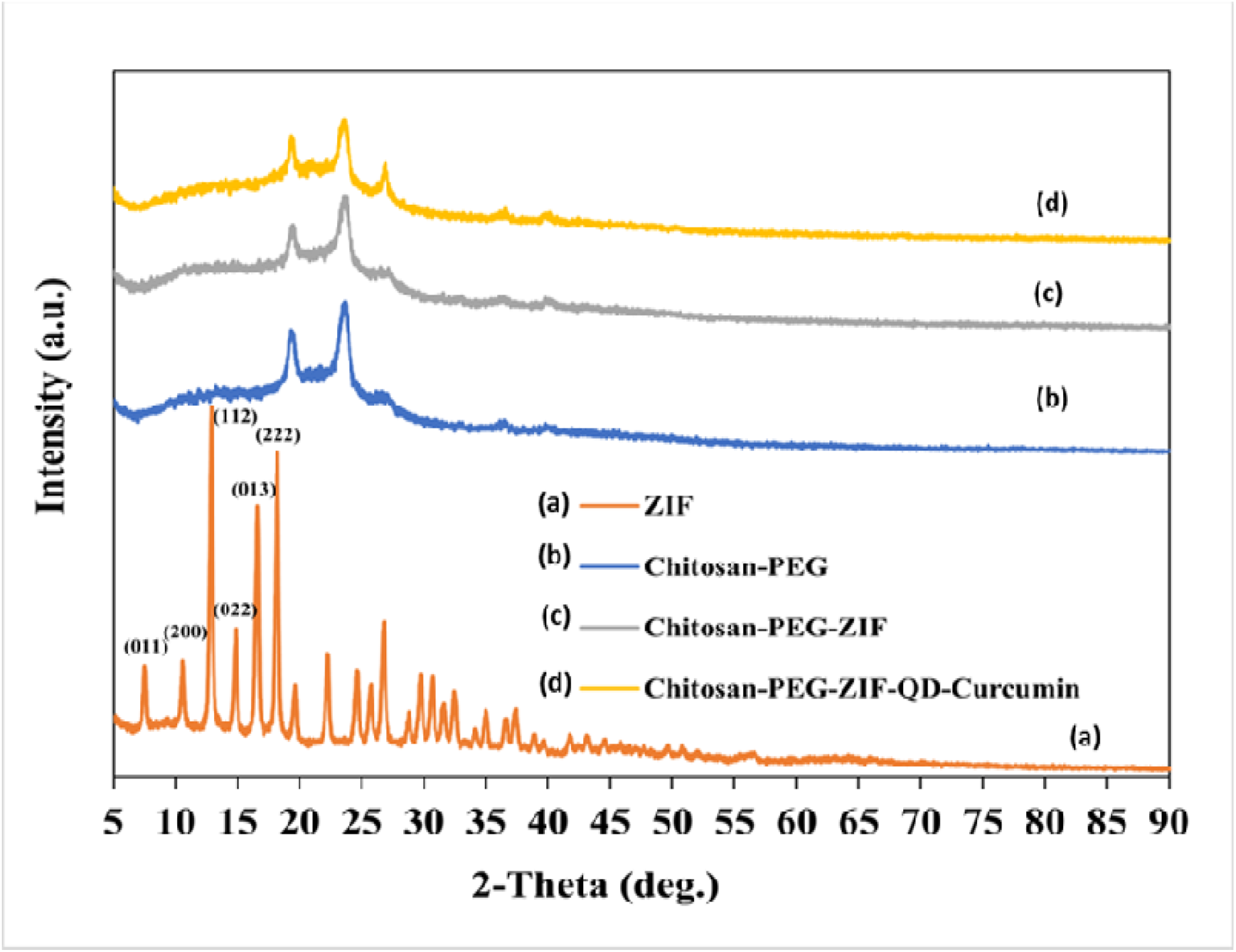
XRD of ZIF(a), CH-PEG (b), CH/PEG/ZIF (c), CH/PEG/ZIF/NCQD/CU (d).

### 3.3. Surface morphology

It is evident from the SEM pictures in Fig. 3 (a, b) that ZIF-90 has a rectangular solid morphology that is almost uniform, exhibiting a smaller than 100 nm size distribution. The prepared samples’ phase purity was validated by SEM pictures. The synthesis process and environmental factors, including the solvent and temperature employed^76^, affect the shape and dimensions of ZIF-90. A key factor influencing the drug release behaviour of a DDS is its surface. Fig.4 (a-d) displays the morphology of the loaded hydrogel nanocomposite. A lamellar structure is clearly visible, with a random distribution of spherically shaped nanocarriers appropriate for drug delivery applications. The core@shell form of the nanocomposite, as seen in Fig. 4(d), was produced using the water-oil-water double emulsion process. The Chitosan/PEG copolymer matrix and sorbitan monooleate with water molecules encircle the drug-loaded ZIF in the core of this core@shell structure. The shell layer may give the nanocomposite stimuli-responsive behaviour and reduce the toxic behaviour of medications and nanoparticles embedded inside the core^77–79^. The addition of sorbitan monooleate stimulates the development of minute droplets, which are equivalent to composite nanospheres. The shape can offer excellent stability, controlled release, and strong drug protection.

**Fig. 3.**
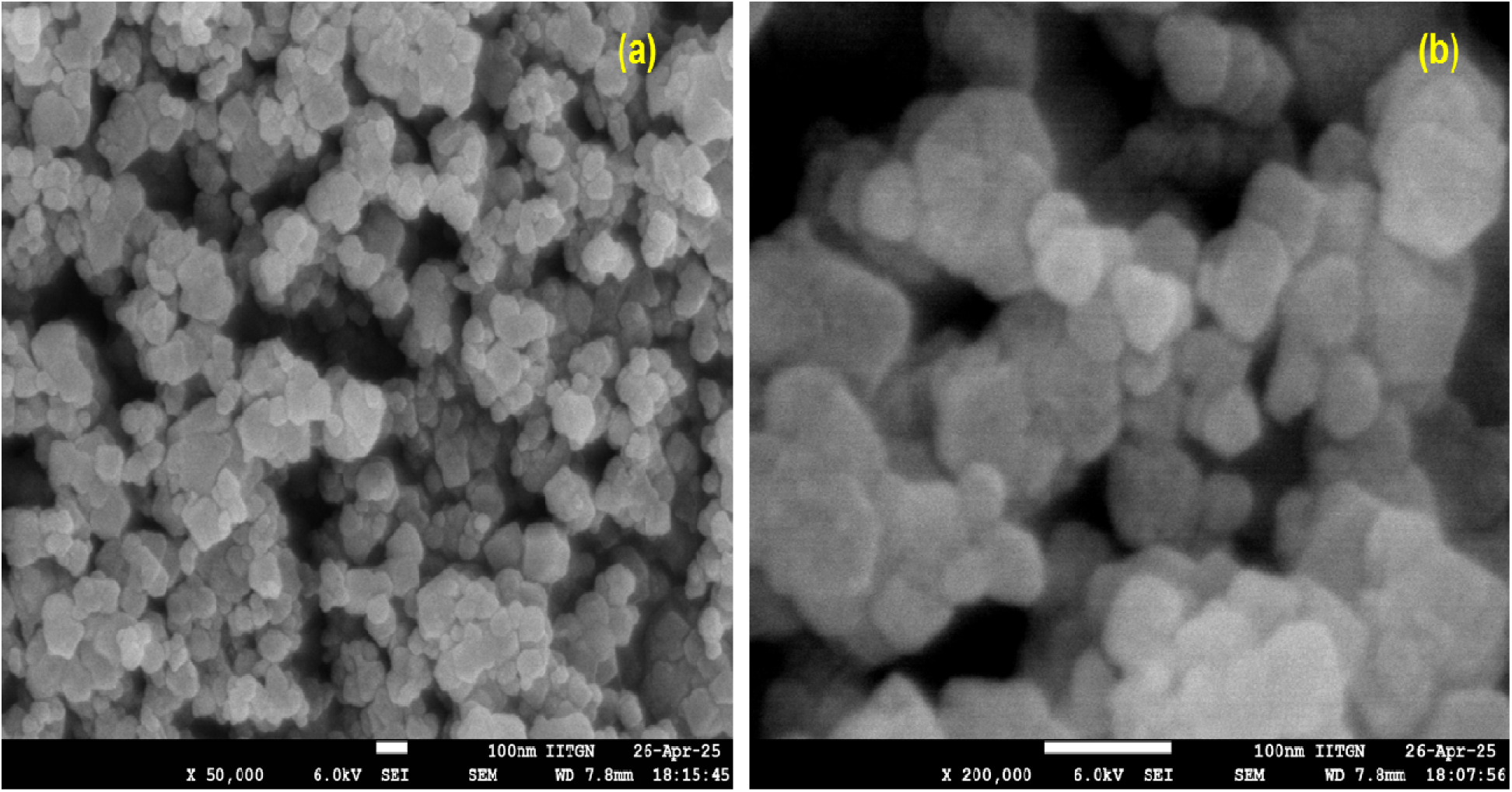
FESEM image of ZIF-90.

**Fig. 4.**
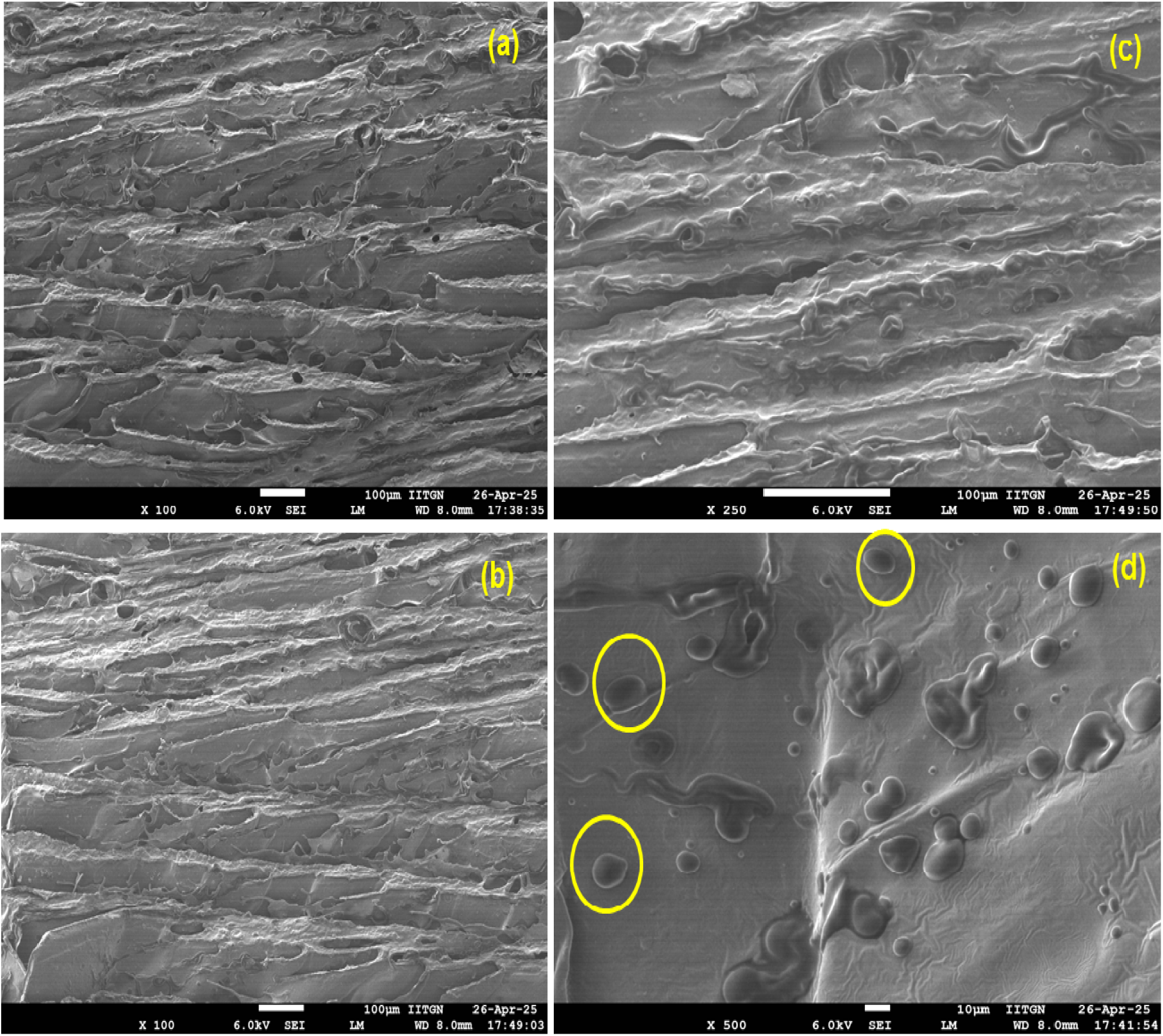
FESEM images of Chitosan/PEG/NCQD/ZIF containing Curcumin.

### 4.4 DLS and Zeta potential

The DLS analysis was carried out to determine the average hydrodynamic radii, as shown in Fig.5. It was determined that the mean particle radius was 150 nm, which is suitable for a nanomedicine delivery method. This outcome validates the NPs’ uniform mixing. The nanocarriers’ tiny particle sizes allowed them to aggregate in tumour locations and take advantage of the EPR effect. NPs < 200 nm were successfully and easily dispersed within capillaries. The diameter of the nano drug varies between 110 and 220 nm, having an overall size of 150 nm. Additionally, the size distribution data shows a single peak, indicating a uniform and monodispersed nano system. Furthermore, Fig. 5 displays an average surface potential of nanocomposites to be approximately +45 mV, which gives them excellent stability in aqueous solutions and the bloodstream. The chitosan amino groups that cover the other nanoparticles in the intended DDS are the cause of this potential. In actuality, nanocarriers cannot assemble due to high repulsive electrostatic forces. It is noteworthy that positive zeta potential may enhance the internalisation of cancerous cells, as demonstrated in the literature^80^. The QD, measuring less than 1 nm in Fig. 6, having a zeta potential of -5 mV, shows no aggregation, which improves the stability of the nanodrug.

**Fig. 5.**
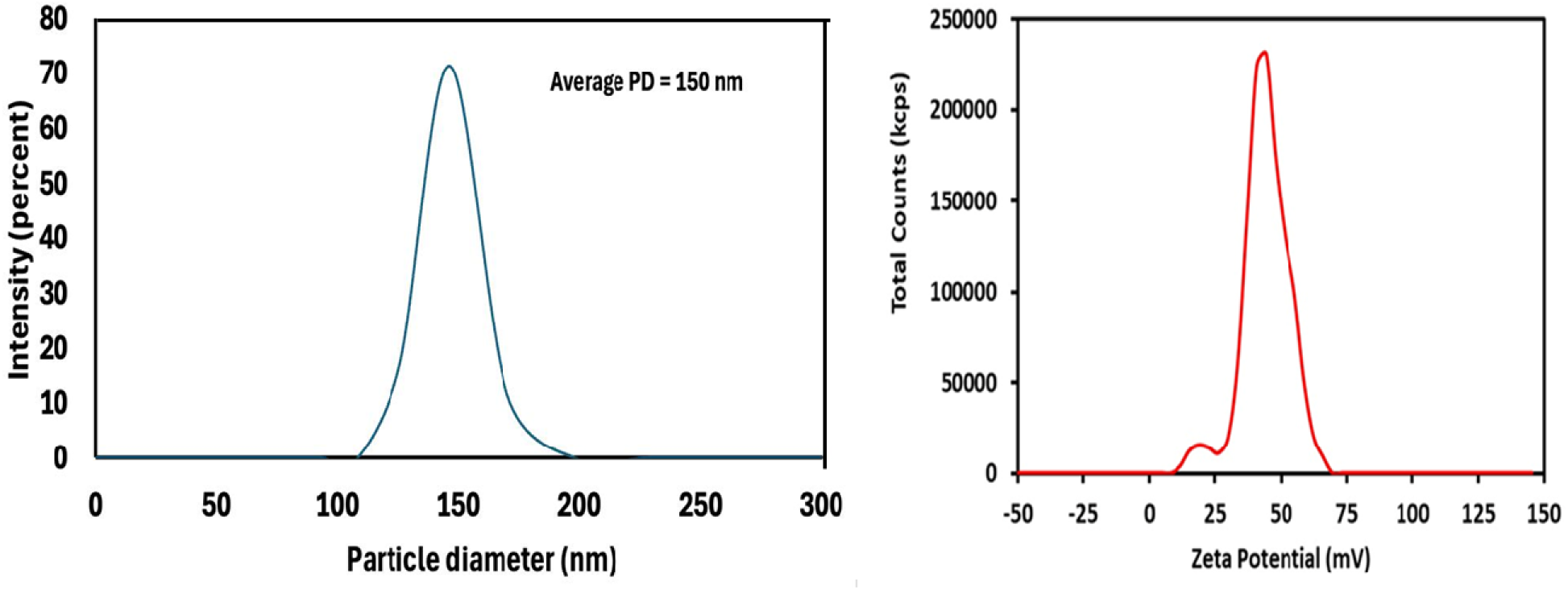
DLS and Zeta Potential of the final nanocomposite with curcumin.

**Fig. 6.**
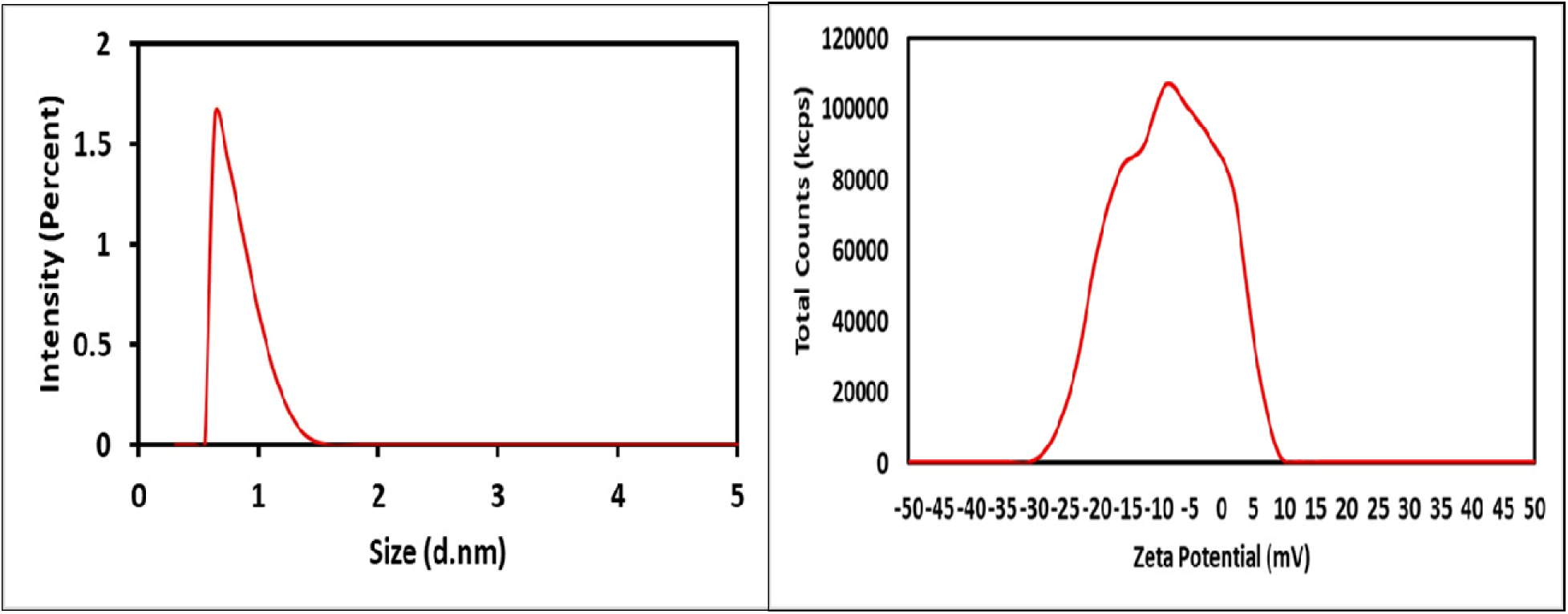
DLS and Zeta potential of NQD.

### 3.5 Optical properties

The UV-visible absorption spectrum (Fig. 7) provides supplementary validation of the composite assembly. In the top spectra of Fig.7, when curcumin is incorporated into a composite material, the absorption maximum, at 425 nm, typically remains unchanged compared to free curcumin. This suggests that the chemical structure responsible for the absorption (the π-π* transition) is not significantly altered by the composite formation. In the bottom spectra of Fig.7, the transition and the transition of in CDs are responsible for the two distinct absorption peaks of NCQDs are seen at 230 and 330 , respectively. The curcumin-loaded NCQDs exhibited an absorption peak at 330 nm and another peak at 425 nm, indicating no shift in the absorption maximum compared to the original carbon quantum dots, except for some broadening on adding higher amounts of curcumin.

**Fig. 7.**
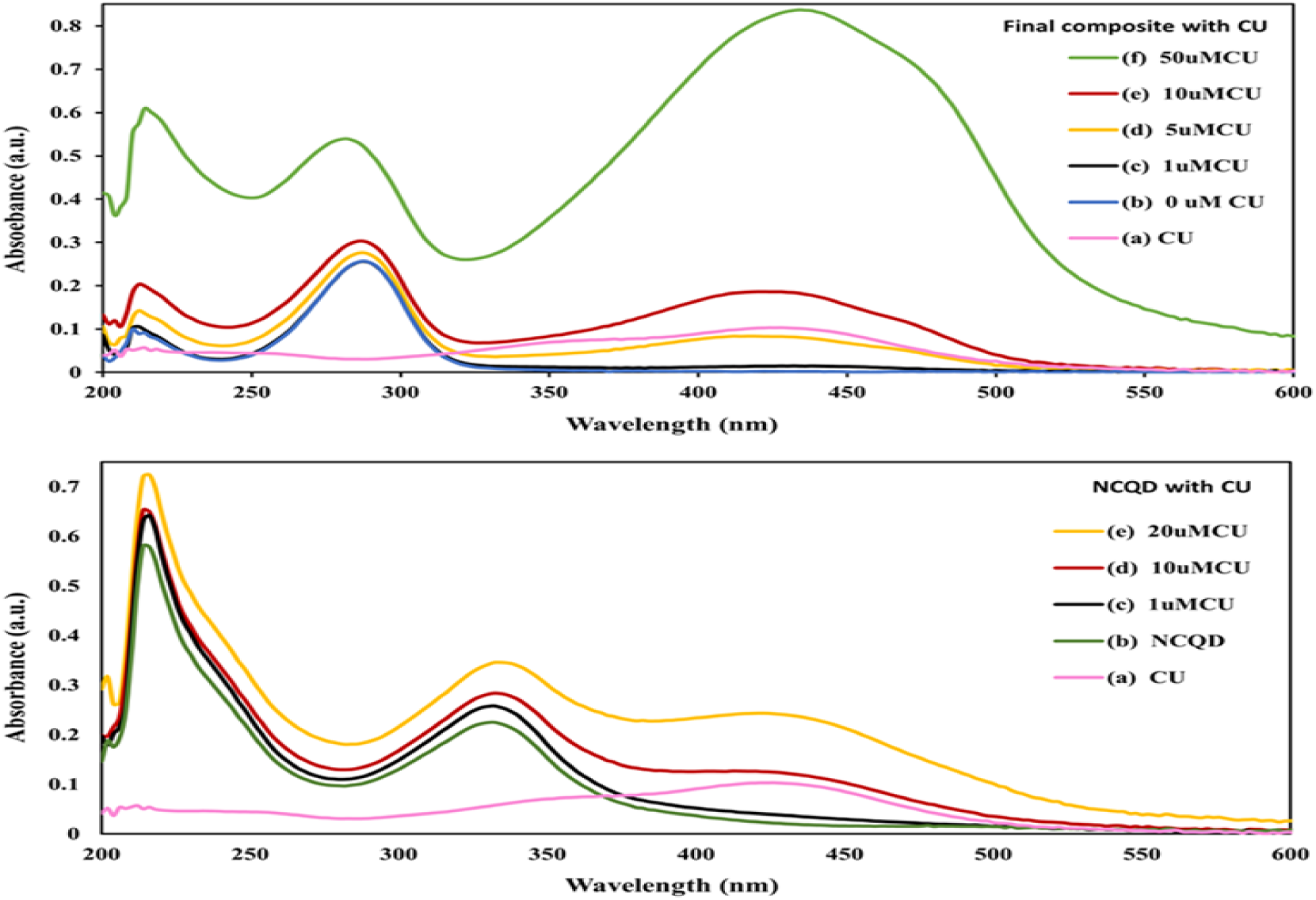
Top: UV spectra of absorbance of Curcumin(a), composite (b), composite with variable conc of curcumin from 1 to 50 µ M (c-f); Bottom: UV spectra of absorbance of CU(a), NCQD (b), NCQD containing different concentrations of curcumin (1 to 20 µ M)

The top spectra in Fig. 8 depict a red shift in the PL spectra of QD from 425 to 530 with the excitation wavelength intensifying from 330 to 410 . Conversely, the PL intensity exhibits clear excitation-dependent characteristics, with a maximum intensity peak at an excitation wavelength of 330 nm. However, the fluorescence intensity in bottom spectra of Fig.8, shows a dramatic drop after adding curcumin due to quenching, where NCQDs are energy benefactors and curcumin acts as an energy absorber. The inset shows the quenching effect of curcumin when added to quantum dots, where QDs are energy providers and curcumin acts as an energy absorber. The most probable quenching mechanism, as seen in Fig. 9, shows the possibility of Förster resonance energy transfer (FRET) validated by an overlap between the emission band of NCQD & the absorption band of curcumin^81^.

**Fig. 8.**
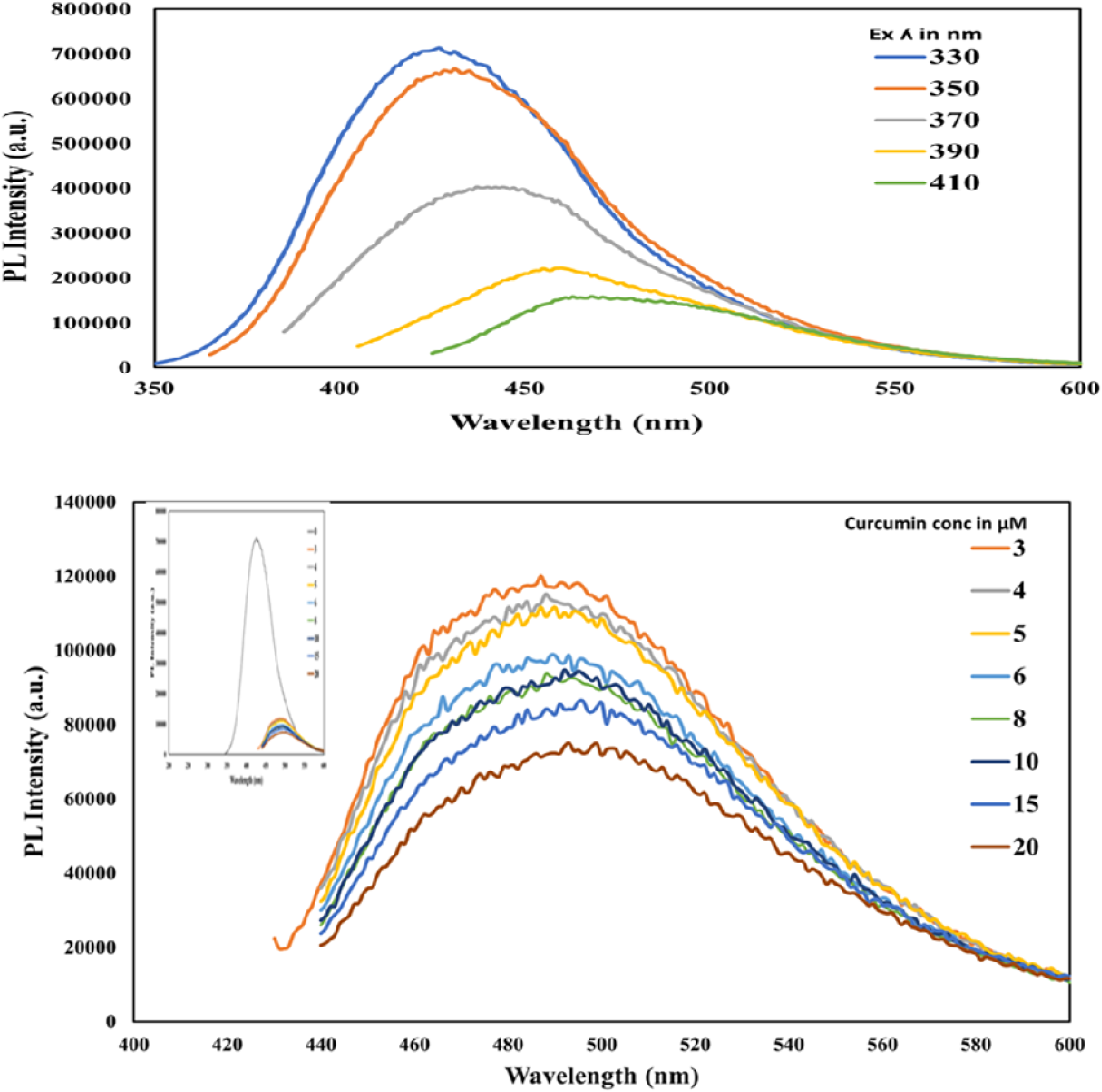
Top: Variation in PL intensity of NCQD. Top: with excitation wavelength varying from 330-410 nm; Bottom: with increasing concentration of curcumin (3 to 20 µM), Inset shows the quenching effect of adding curcumin.

**Fig. 9.**
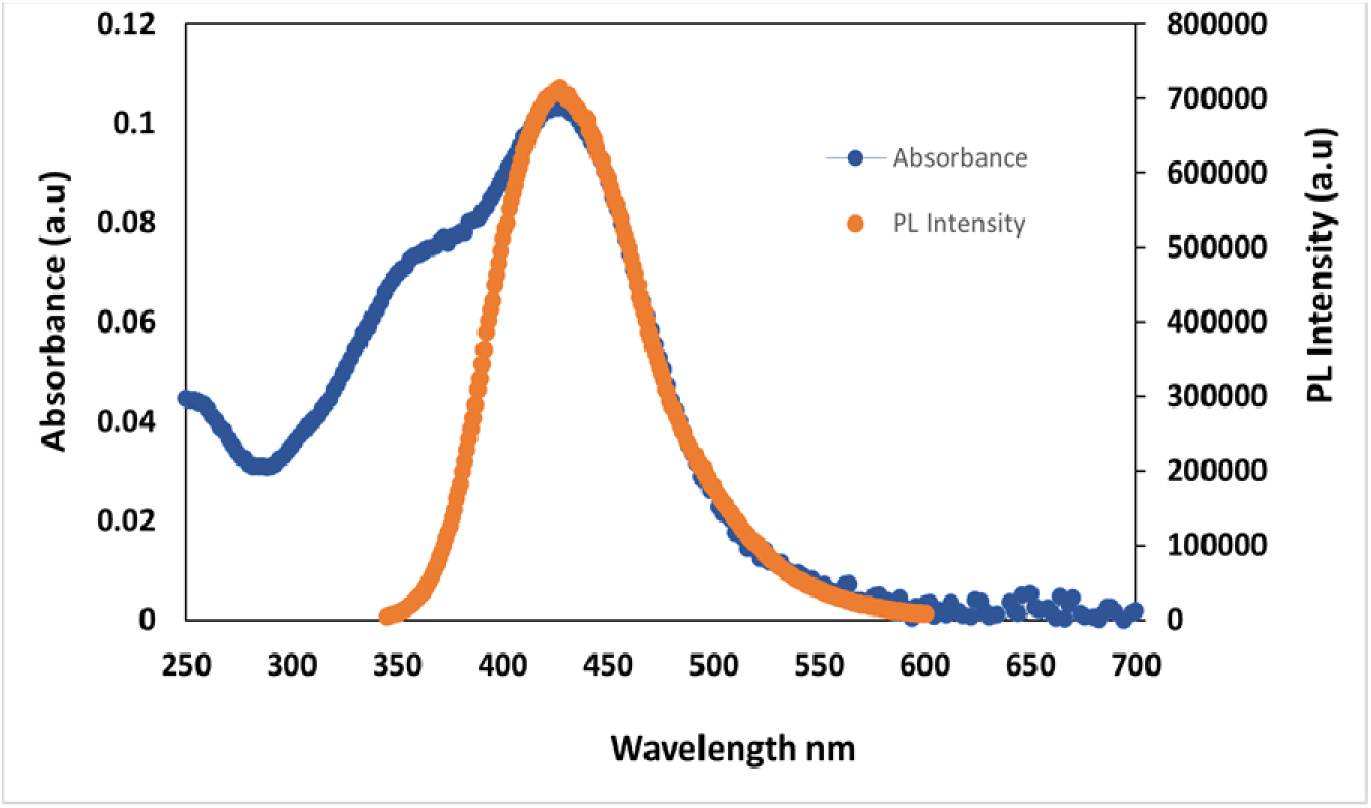
Quenching showing FRET between NCQD & Curcumin.

### 3.5 Curcumin release profile

An initial burst release is observed for the first 6 h until the release percentage reaches 11 %, followed by a gradual and more sustained release. After 12 h, the percentage of drug released in an acidic environment is considerably higher than in a neutral environment.

The transport of drug from nanocarriers at acidic (5.4) and neutral pH (7.4) buffer medium at 37 °C was evaluated using the dialysis procedure over 96 hours (Fig. 10). Because of the links between the NPs, the modelled drug, and the polymers, the total discharge of curcumin at neutral pH is lower than at acidic pH, maintaining the integrity of the nanocomposite. The nanocarrier carries the medication to the targeted tumour cells, and controlled-release behaviour is noted when the DDS is near the healthy cells.

**Fig. 10.**
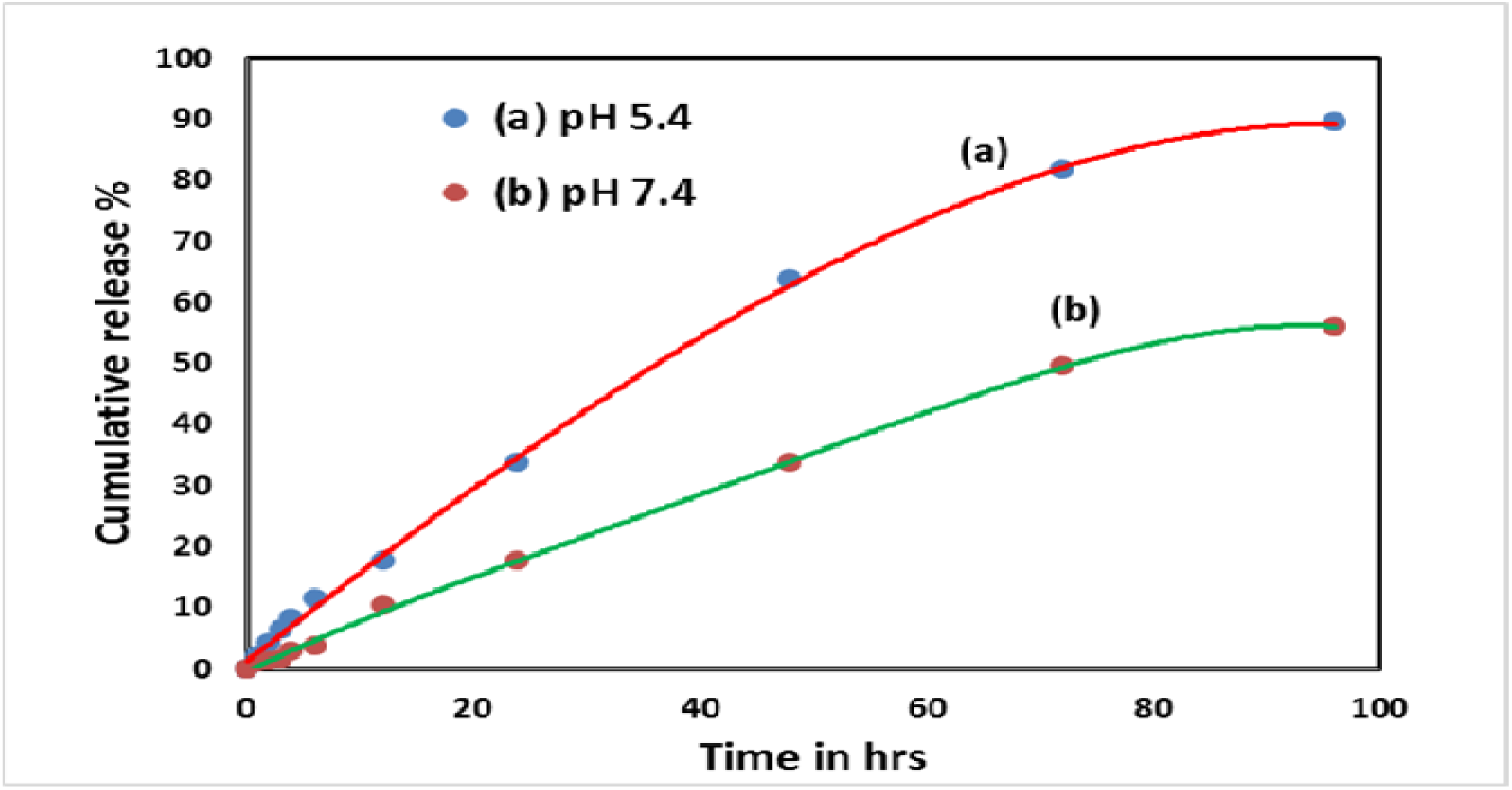
Curcumin release profile of the final DDS at pH 5.4 and 7.4.

The alteration in the release mechanism at different pH levels demonstrates the nanocarrier’s pH responsiveness. In acidic conditions, ZIF is stimulated to break down by protonation. A repulsive contact amongst the neighbouring +ve charge in Zn^2+^ and other protonated species, such as carbonyls, hydroxyls, and amines of other DDS constituents, facilitates the disintegration of the nanostructure in acidic conditions. This increases the amount of drug delivery.

Figs. 11 shows the drug release data fitted on zero and first-order, Higuchi, and Korsmeyer-Peppas (K-P) kinetic models. Parameters such as R^2^ values and kinetic characteristics, as investigated, are provided in Table 2. The highest R^2^ values are equal to 0.998 at 7.4 for the zero order model, and 0.996 for the K-P model at 5.4 pH, respectively. Zero-order release kinetics refers to the process of drug release at a constant rate from a drug delivery system (i.e., the same amount of drug is released per unit time independently of the amount of drug remaining in the delivery system. The Korsmeyer-Peppas model is a mathematical framework used to describe drug release kinetics from polymeric systems, particularly those with controlled release mechanisms.

**Fig. 11.**
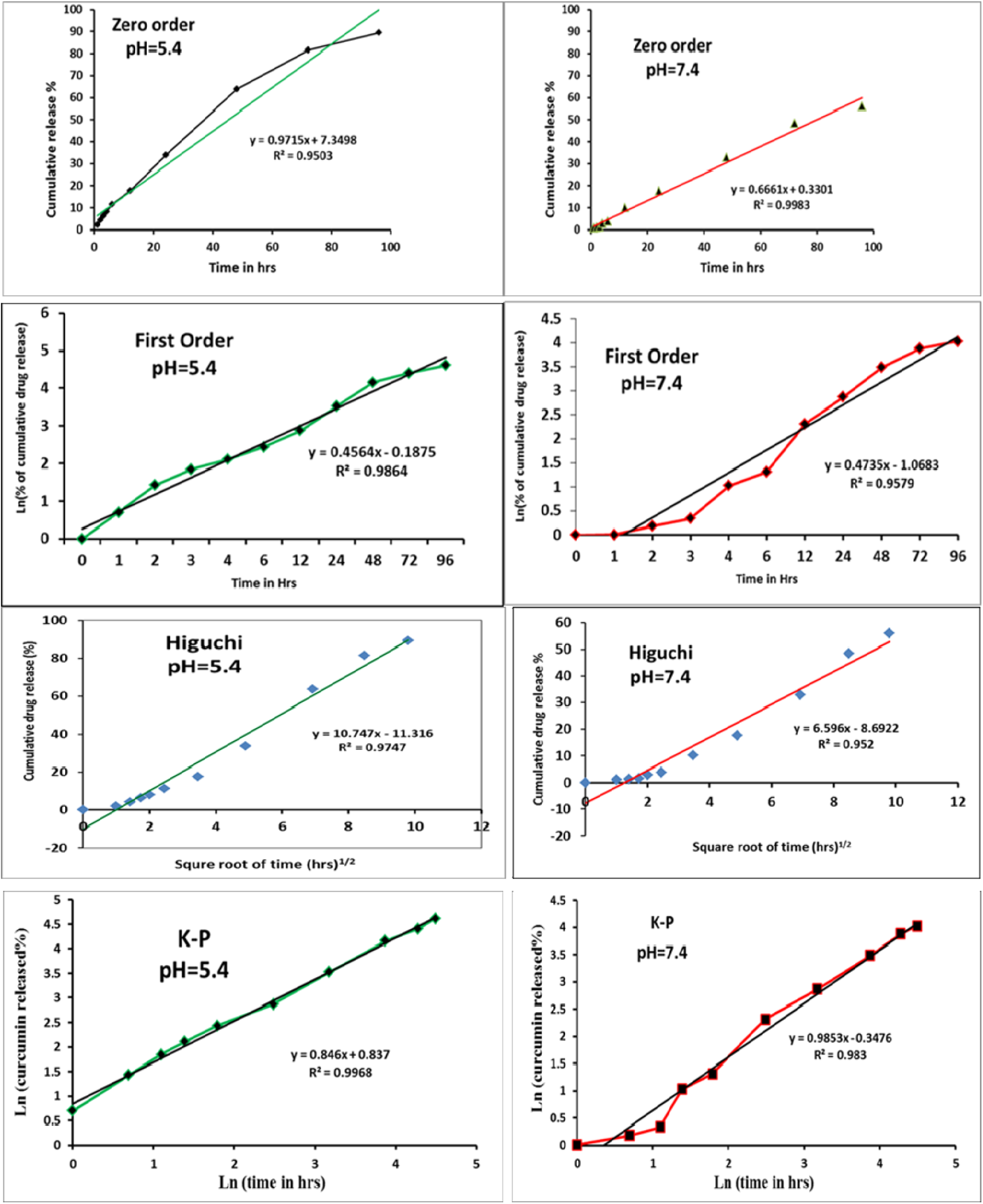
Different kinetic models for curcumin release from the composite at pH 5.4 and 7.4.

**Table. 1:**
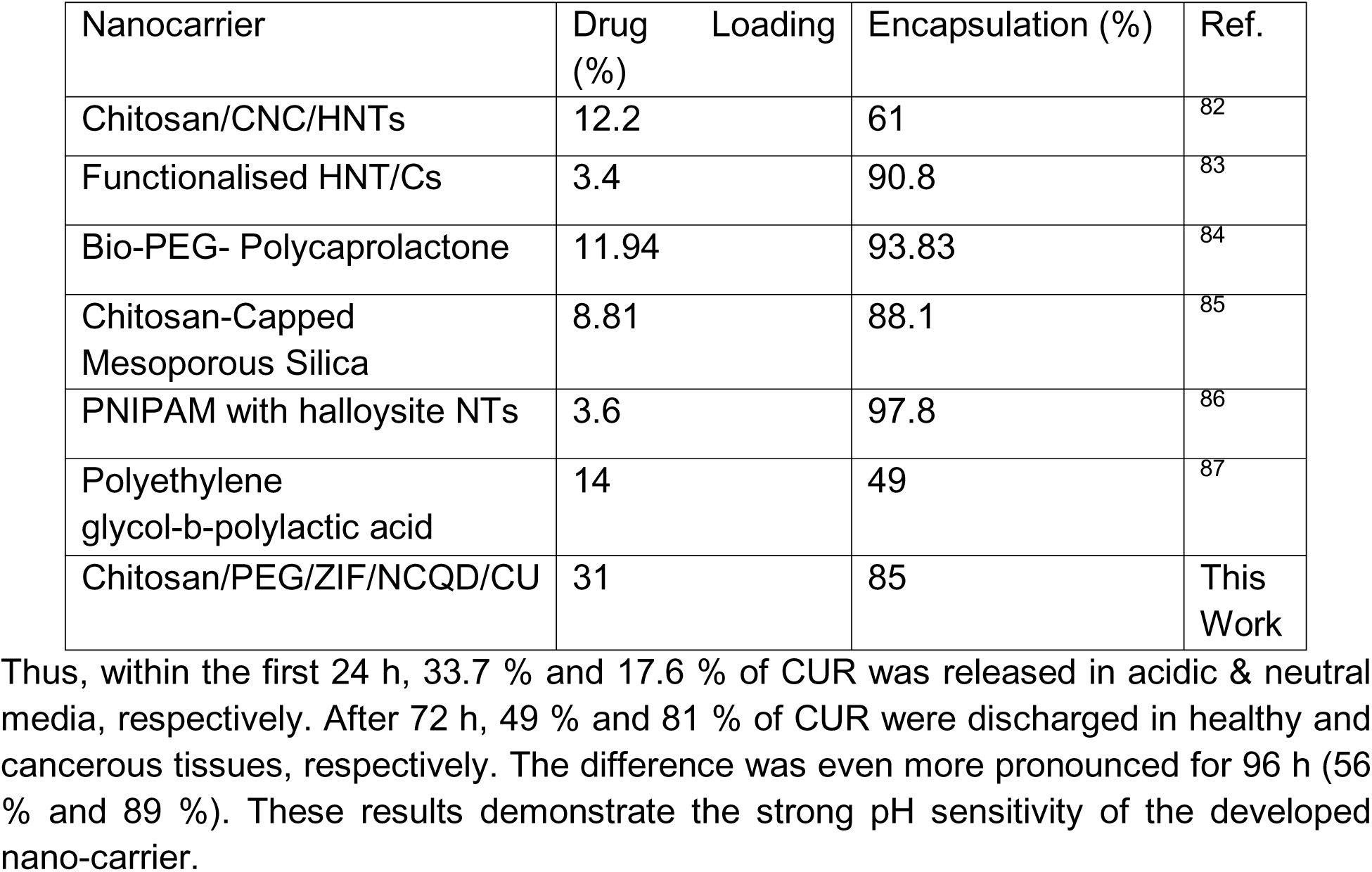
A comparative data of curcumin encapsulation and loading efficiencies.

**Table 2:**
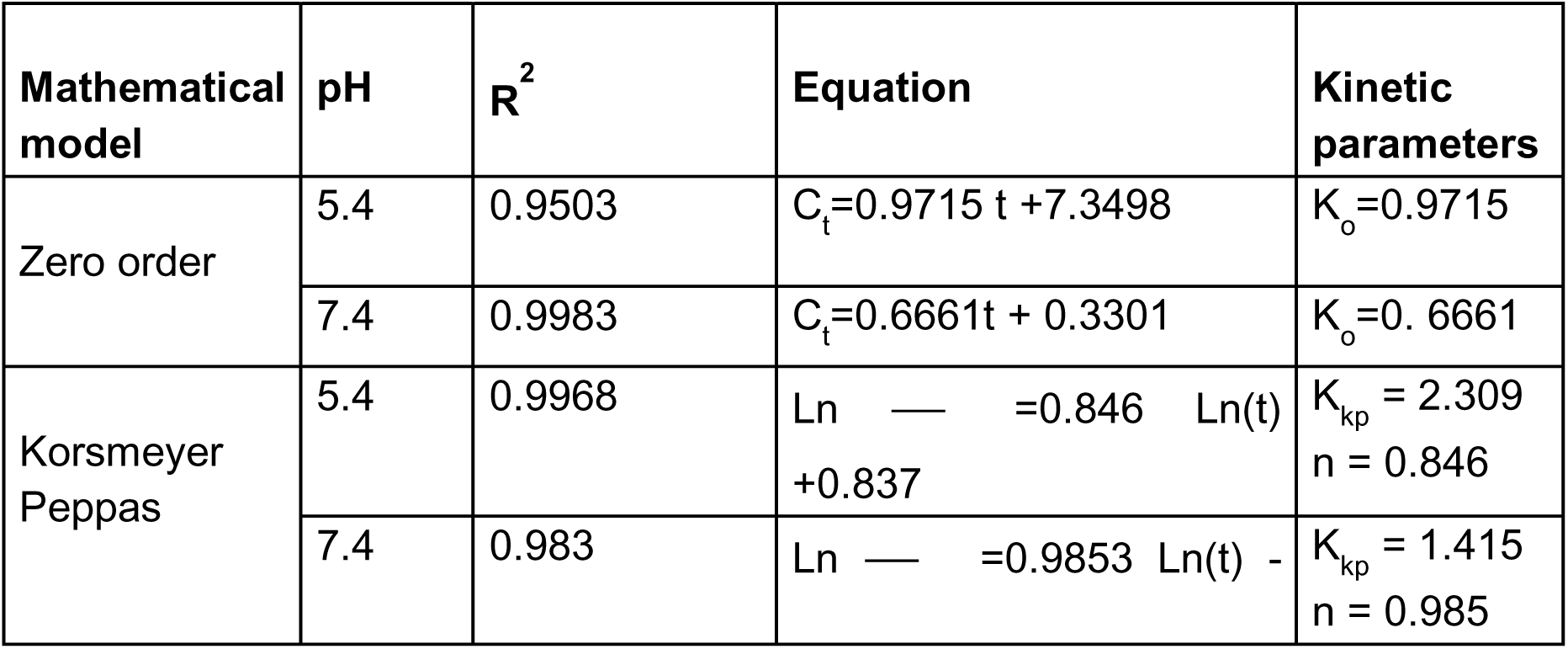
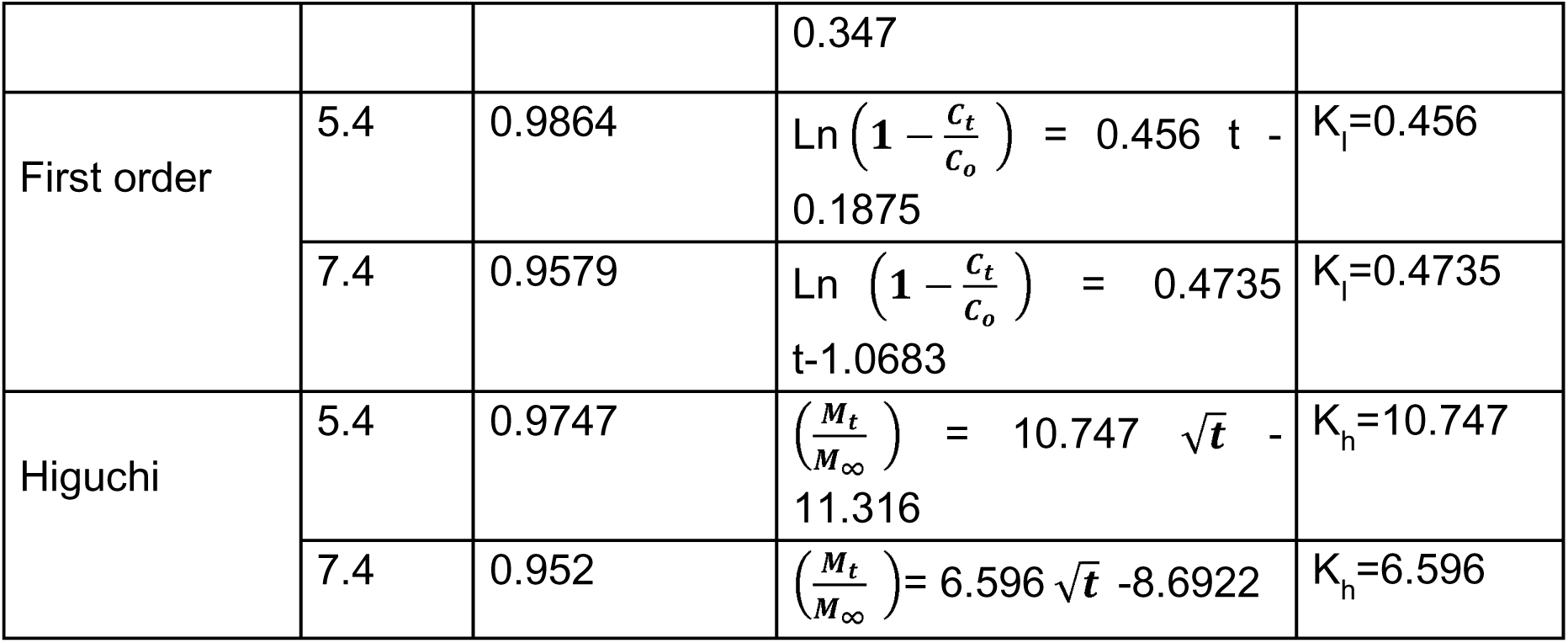
Different mathematical models for drug release kinetic analysis.

### 3.6 MTT Assay

Fig. 12 displays the bar graph with the computed percentage of cell viability in relation to the control group. The anticancer activity of the composite emulsion was assessed 72 hours after cell treatment in line with kinetic study. Using the IC_50_ Probit technique, the IC50 value of composite emulsion was calculated after 72 hours of treatment on MCF-7. The curcumin-loaded Chitosan/PEG/NCQD/ZIF emulsion demonstrated an IC_50_ of 17.5 μg/mL and the cell viability significantly decreased at all tested concentrations, indicating a strong anticancer impact (**** p<0.0001). According to the literature^4,88–90^, the IC_50_ of free curcumin against MCF-7 breast cancer cells usually varies between 10 and 25 μM, depending on the particular experimental setup. In summary, our experimental data strongly support the potential of the synthesized pH-sensitive nanocarrier for controlled delivery of anticancer medications.

**Fig. 12.**
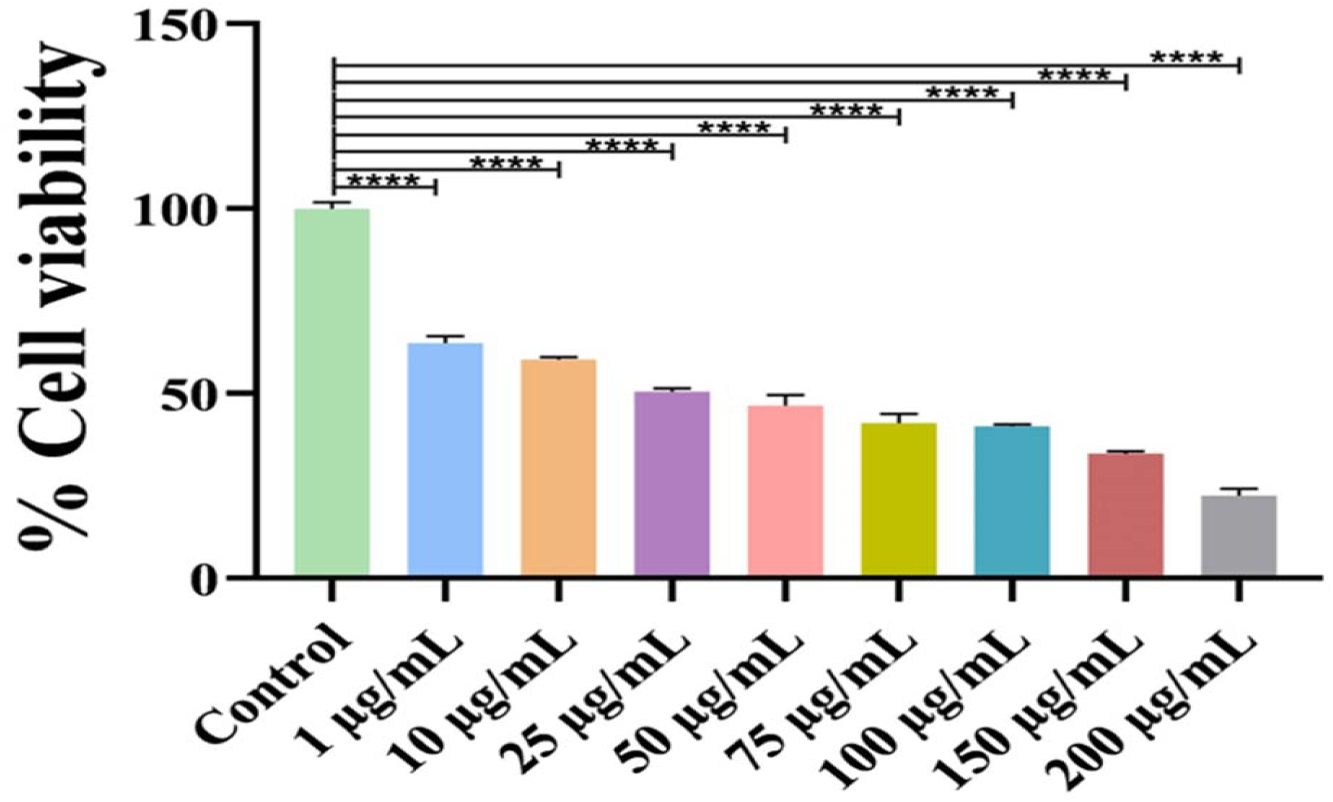
In vitro cytotoxicity of final composite in comparison to control group (**** denotes p<0.0001)

## Conclusions

NP-based drug delivery systems are linked to better pharmacokinetics, biocompatibility, tumor targeting, and stability when compared to conventional medications. In this study, ZIF was added to a drug delivery system to enhance its ability to load curcumin, and its efficacy was tested against breast cancer cells. XRD, FTIR, and DLS measurements were used to examine the manufactured system’s physicochemical characteristics. DLS confirmed the FESEM’s good dispersion of appropriately sized nanoparticles with an average particle size of 150 nm. This allows nanocomposites to penetrate tissues and target particular body locations. Additionally, the amide functional groups produced a positive 54 mV surface charge in the zeta potential finding, indicating a stable platform in blood. The Chitosan/PEG network’s loading (31%) and entrapment efficacy (85%) were greatly enhanced by adding ZIF and QDs.

By comparing the drug release profiles of NDDS with varying pH, it was observed that this NDDS is pH-sensitive and capable of targeting malignant cells with 89% release after 96 hrs compared to 56% release at normal pH. The drug released per unit time at 7.4 pH, following zero-order kinetics, was independent of the amount of drug left in the delivery system. In contrast, the Korsmeyer-Peppas model verified drug release kinetics with a controlled release mechanism at pH 5.4. Further research is necessary to evaluate the therapeutic efficacy of the MOF-based composite drug delivery system, although the results indicate that it may find application in clinical settings.

## Future Perspective

In this work, a new pH-sensitive nanocarrier platform is suggested by using MOF-composites with carbon quantum dots (CQDs) and a chitosan/PEG-based stabilizing matrix. This platform significantly enhances the solubility, bioavailability, and controlled release characteristics of curcumin, thus explaining its high potential to be used in targeted cancer therapy. In addition, the addition of DNA nanotechnology to this composite system has exciting potential towards the creation of the next generation of smart therapeutics. The DNA nanostructures, including DNA tetrahedra and DNA origami, are highly programmable, biocompatible, and structurally precise, which makes them perfectly applicable to the engineering of stimulus-responsive systems^14,91–99^. These motifs could be included in the MOF-CQD scaffold to allow improved cellular uptake, controlled disassembly in response to tumor-specific cues, and even molecular logic-gated drug release.

The carbon quantum dots (CQDs) have photoluminescence and biocompatibility, which make them passive fluorescent markers, and, simultaneously, allow them to be used as active elements in real-time drug tracking and diagnostic sensing^100–102^. CQDs may be integrated into Förster resonance energy transfer (FRET) systems by proper surface modification to detect drug release kinetics or intracellular interactions. Moreover, the nanocomposite surface can be functionalized by DNA aptamers or antisense RNA strands to increase tumor specificity and minimize off-target effects. The DNA moiety in this arrangement acts as a targeting ligand as well as a switch.

The co-loading capability of metal-organic frameworks (MOFs) and DNA-based nanostructures allows the creation of multimodal carriers that can also integrate several therapeutic agents. These platforms can deliver combinatorial therapeutic regimens with temporal or spatial control release profiles and thus allow precision therapy. They have a modular architecture which enables the inclusion of various functional modules which are triggered by different stimuli, especially pH, redox conditions and enzymatic activity, which makes them especially applicable to heterogeneous tumor microenvironments. Despite the feasibility and in vitro biocompatibility shown by the present study, future research directions must be focused on biodistribution, pharmacokinetics, and immunological responses in vivo. Increasing the synthesis approach to guarantee batch reproducibility and regulatory alignment will also be critical to the advancement toward clinical translation.

The current research lays the basis of the future generation of multifunctional, trackable, and highly specific nanotherapeutics. Using the synergistic strengths of metalorganic frameworks (MOFs), carbon quantum dots (CQDs), and DNA nanotechnology, future iterations of this platform may become smart drug delivery platforms that simultaneously combine therapeutic, diagnostic, and real-time feedback functions into a single nano-scale structure.

## Acknowledgements

We thank ANRF-CRG, GSBTM, IITGN and MoES-STARS for research funding. We declare no conflict of interest.

